# Humans can visually judge grasp quality and refine their judgments through visual and haptic feedback

**DOI:** 10.1101/2020.08.11.246173

**Authors:** Guido Maiello, Marcel Schepko, Lina K. Klein, Vivian C. Paulun, Roland W. Fleming

## Abstract

How humans visually select where to grasp objects is determined by the physical object properties (e.g., size, shape, weight), the degrees of freedom of the arm and hand, as well as the task to be performed. We recently demonstrated that human grasps are near-optimal with respect to a weighted combination of different cost functions that make grasps uncomfortable, unstable or impossible e.g., due to unnatural grasp apertures or large torques. Here, we ask whether humans can consciously access these rules. We test if humans can explicitly judge grasp quality derived from rules regarding grasp size, orientation, torque, and visibility. More specifically, we test if grasp quality can be inferred (i) by using motor imagery alone, (ii) from watching grasps executed by others, and (iii) through performing grasps, i.e. receiving visual, proprioceptive and haptic feedback. Stimuli were novel objects made of 10 cubes of brass and wood (side length 2.5 cm) in various configurations. On each object, one near-optimal and one sub-optimal grasp were selected based on one cost function (e.g. torque), while the other constraints (grasp size, orientation, and visibility) were kept approximately constant or counterbalanced. Participants were visually cued to the location of the selected grasps on each object and verbally reported which of the two grasps was best. Across three experiments, participants could either (i) passively view the static objects, (ii) passively view videos of other participants grasping the objects, or (iii) actively grasp the objects themselves. Our results show that participants could already judge grasp optimality from simply viewing the objects, but were significantly better in the video and grasping session. These findings suggest that humans can determine grasp quality even without performing the grasp—perhaps through motor imagery—and can further refine their understanding of how to correctly grasp an object through sensorimotor feedback but also by passively viewing others grasp objects.

## 1 Introduction

When we try to grasp objects, within out field of view we rarely fail. We almost never miss the object or have it slip out of our hands. Thus, humans can very effectively use their sense of sight to select where and how to grasp objects. Yet for any given object, there are numerous ways to place our digits on the surface. Consider a simple sphere of 10 cm diameter and ~300 cm^2^ surface area. If we coarsely sample the surface in regions of 3 cm^2^, (a generous estimate of the surface of a fingertip) there are approximately 100 surface locations on which to place our digits. Even when considering simple two-digit precision grips, which employ only the thumb and forefinger, there are ~10,000 possible digit configurations that could be attempted. How do humans visually select which of these configurations is possible and will lead to a stable grasp?

To answer this question, in recent work (Klein, Maiello et al., 2020) we asked participants to grasp 3D polycube objects made of different materials (wood and brass) using a precision grip. Even with these objects—decidedly more complex than a simple sphere—participants consistently selected only a handful of grasp configurations, with different participants selecting very similar grasps. This suggests that a common set of rules constrains how people visually select where to grasp objects. We formalized this observation, following (Kleinholdermann et al., 2013), by constructing a computational model that takes as input the physical stimuli, and outputs optimal grasp locations on the surface of the objects. Specifically, we constructed a set of optimality functions related to the size, shape, and degrees of freedom of the human hand, as well as to how easily an object can be manipulated after having been grasped. Model predictions closely agreed with human data, demonstrating that actors choose near-optimal grasp locations following this set of rules.

The strongest constraint for two-digit grasps, included in this computational framework, requires surface normals at contact locations to be approximately aligned (a concept known as force closure; Nguyen, 1988). Fingertip configurations that do not fulfill this constraint, e.g. with thumb and forefinger pushing on the same side of an object, cannot lift and manipulate the object. Indeed successful human grasps never fail to meet the force closure constraint (Klein, Maiello et al., 2020; Kleinholdermann et al., 2013). The other constraints we implemented as optimality functions relate to:

> *Natural grasp axis*: humans exhibit a preferred hand orientation for precision grip grasping, known as the natural grasp axis (Lederman & Wing, 2003; Roby-Brami et al., 2000; Schot et al., 2010; Voudouris et al., 2010), which falls within the midrange of possible hand and arm joint angles. Grasps rotated away from the natural grasp axis may result in uncomfortable (or impossible) hand/arm configurations that require extreme joint angles. Since these extreme joint angles should be avoided (Rosenbaum et al., 2001), optimal grasps should exhibit minimum misalignment with the natural grasp axis.
>
> *Grasp aperture*: When free to employ any multi-digit grasp, participants select precision grip grasps only when the required distance between finger and thumb at contact (the ‘grasp aperture’) is smaller than 2.5 cm (Cesari & Newell, 1999). As grasp size increases, humans progressively increase the number of digits employed in a grasp. Therefore, optimal two-digit precision grips should exhibit grasp apertures below 2.5 cm.
>
> *Minimum torque*: grasping an object far from its center of mass results in high torques, which may cause the object to rotate when manipulated (Eastough & Edwards, 2006; Goodale et al., 1994; Lederman & Wing, 2003; J. Lukos et al., 2007; Paulun et al., 2016). Large gripping forces would be required to counteract high torques and prevent the object from rotating. Thus, optimal grasps should have minimum torque.
>
> *Object visibility*: when grasping an object, the hand might occlude part of an object from view. This could be detrimental for subsequent object manipulation, and indeed humans exhibit spatial biases in their grasping behavior which are consistent with avoiding object occlusions (Maiello, Paulun et al., 2019; Paulun et al., 2014). Therefore, optimal grasps should minimize the portion of an object occluded from view.

Whereas the force closure constraint is necessary and immutable, the relative importance given to the other four constraints varies with object properties (e.g. mass) and across participants.

Given the computational costs, it seems relatively unlikely that the brain fully computes these optimality functions for every possible grasp. Nevertheless, our previous findings suggest that humans can employ visual information to estimate these constraints and guide grasp selection. As a further test of our framework for understanding human grasp selection, here we ask whether human participants can explicitly report relative grasp optimality (i.e., which of two candidate grasps would be closer to optimal). We further ask whether observers can judge grasp optimality using vision alone, or whether executing a grasp is necessary to do so.

If participants were indeed better at judging grasp optimality when executing grasps, this might suggest that tactile (Johansson & Westling, 1984) and proprioceptive feedback from our arm and hand (Lukos et al., 2013; Rosenbaum et al., 2001) plays a role in evaluating grasp quality. Humans may employ these sources of feedback to learn that certain hand configurations are uncomfortable, or that one grasp requires more force than another to pick up the same object.

Additionally, participants might also be able to visually assess the characteristics of the own movements, such as the speed and trajectory of the limb. These sources of visual information are known to play a strong role in grasp planning and execution, as removing them changes the kinematics of grasping movements (Connolly & Goodale, 1999), and even simply observing others execute grasping tasks can improve one’s own grasping performance (Buckingham et al., 2014). We therefore ask how much these sources of visual information might contribute to participant judgements of grasp optimality. Specifically, we test whether grasp quality can be inferred from watching grasps executed by others. If this were the case, then perhaps vision and proprioception may be redundant sources of information about grasp quality, which could aid humans in linking vision and motor control in action planning.

To test whether humans can explicitly judge grasp quality, in Experiment 1 we asked participants to report which of two candidate grasps on an object is best, first using vision alone (vision session), and then also by attempting both grasps on the object, one after the other (grasping session). To test whether visual information about the grasping movements plays a role in judging grasp quality, in Experiment 2a we asked a new set of participants to repeat a subset of key conditions from Experiment 1, while we video-recorded their grasping movements. Finally, in Experiment 2b we showed these recorded movements to yet another set of participants (video session), and asked them to judge grasp quality from the videos of grasps executed by participants from Experiment 2a.

## 2 Methods

### 2.1 Participants

We recruited 21 naïve and right-handed participants (16 female, 5 male; mean [range] age: 24 [19 - 32] years) for Experiment 1, 25 naïve and right-handed participants (17 female, 8 male; mean [range] age: 23 [20 - 26]) for Experiment 2a, and 25 naïve and right-handed participants (18 female, 7 male; mean [range] age: 24 [19 - 36]) for Experiment 2b. Participants were staff and students from Justus Liebig University Giessen, Germany. In return for their participation, volunteers were paid 8 EURO per hour. Participants reported healthy upper extremities and normal or corrected to normal vision. All provided written informed consent. All procedures were approved by the local ethics committee of Justus Liebig University Giessen (Lokale Ethik-Kommission des Fachbereichs 06, LEK-FB06; application number: 2018-0003) and adhered to the tenets of the declaration of Helsinki.

### 2.2 Apparatus

All Experiments (1, 2a, 2b) were programmed in Matlab version 2018a. Participants were seated at a table with a mounted chin rest in a brightly lit room. Figure 1 shows a schematic of the setup. In all experiments, during the vision (Figure 1A) and grasping sessions (Figure 1B), subjects positioned their heads in the chinrest before each trial. Stimulus objects were positioned 34 cm in front of the participant. At this predefined position, a turntable allowed the experimenter to precisely set object orientation. The target location was shifted 23 cm to the right side from the initial object location along the horizontal axis, at a distance of 40 cm relative to the participant. The starting position for the right thumb and index finger was 24 cm to the right and 22 cm in front of the participant. In grasping sessions, objects were grasped with a precision grip at two predetermined locations. A ZED Mini stereo camera (Stereolabs) was attached to the front of the forehead rest to record (720p, 30 fps) grasping movements in Experiments 2a and 2b. To record videos, a simple recording program was written in C++, using the ZED SDK, and called from within the Matlab environment. The camera orientation was adjustable along the z-axis and fixed at an angle of 25° to capture the whole movement sequence. During the experiment, participants did not see the camera due to its position right in front of their forehead (Figure 1B). In Experiment 2b (Figure 1C), videos were presented on an Asus VG248QE monitor (24″, resolution = 1920 x 1080 pixel) at 60 Hz, positioned at a distance of 40 cm from the observers.

**Figure 1.**
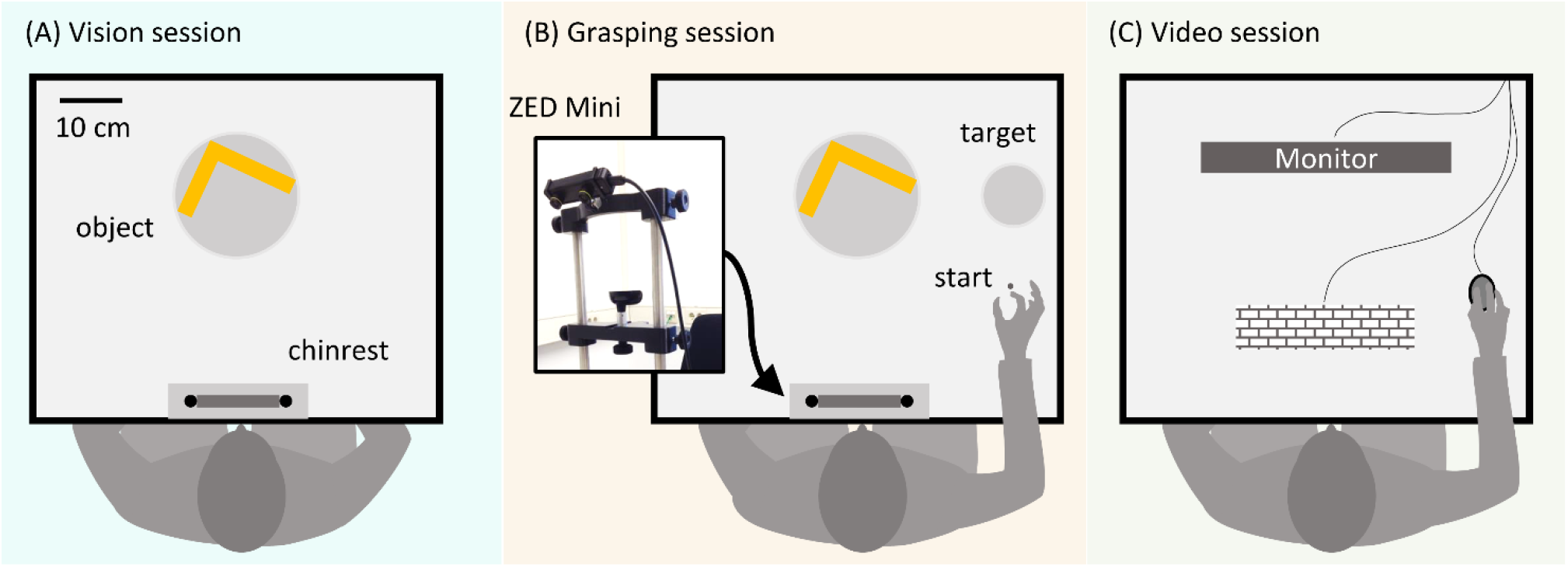
Schematic representation of the experimental setup. **(A)** In the vision sessions participants passively viewed objects and evaluated the relative optimality of preselected grasps without executing the grasps **(B)** In the grasping sessions, participants executed grasps prior to judging the grasp quality. A ZED Mini stereo camera positioned above the participant’s forehead recorded the grasping movements in Experiments 2a and 2b **(C)** In the video session, participants from Experiment 2b viewed recordings of grasps executed by participants from Experiment 2a on a computer monitor.

### 2.3 Experiment 1

#### 2.3.1 Stimuli

In Experiment 1, we employed 16 3D objects (4 shapes, 4 material configurations), each made of 10 cubes (2.53 cm^3^) of beech wood or brass. Objects with the same shape but different material configuration varied in mass (light wooden objects: 97 g, heavy wood/brass objects: 716 g) and mass distribution. These objects were the same, and were presented at the same orientations, as previously described in (Klein, Maiello et al., 2020). For each of the objects, we selected pairs of grasps, one near-optimal and one sub-optimal, according to one of four grasp optimality criteria: natural grasp axis; optimal grasp aperture; minimum torque; optimal visibility. These criteria were mathematically defined as in (Klein, Maiello et al., 2020). For each of these optimality criteria, we selected pairs of near-optimal and sub-optimal grasps on four of the 16 objects, while maintaining the other optimality criteria approximately constant across the grasp pair or counterbalanced across objects. Figure 2A shows one example object in which we selected one near-optimal and one sub-optimal grip with regard to grasp aperture. Figure 2B shows the optimality values for both grasps following each of the optimality criteria, and the difference in optimality between the two grasps. The difference in grasp optimality between pairs of grasps on all 16 objects for each of the four grasp optimality criteria is shown in Figure 2C. The selected grasp pairs were marked on the objects with colored stickers glued onto the objects’ surface. Thumb grasp locations were marked in either blue or green (randomly assigned to the near-optimal and sub-optimal grasps). Index finger locations were marked in yellow. All objects and selected grasp pairs are shown in Supplementary Figures 1-4.

**Figure 2.**
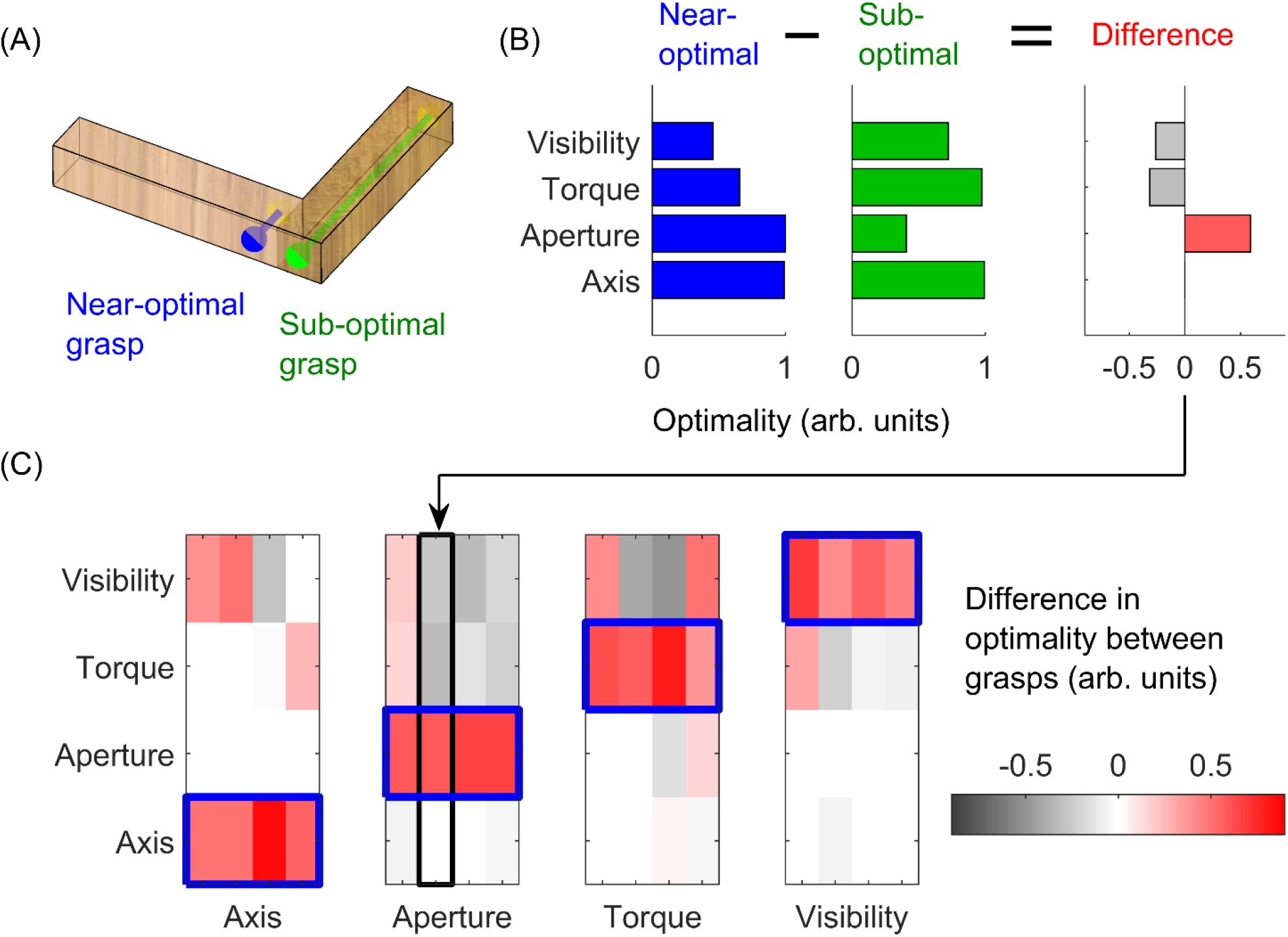
Stimulus selection. **(A)** One example object in which we selected one optimal (blue) and one sub-optimal (green) grasp with respect to grasp aperture. The right side of the object is made of brass, the left side of beech wood. Blue and green dots represent thumb contact locations; the index finger is to be placed on the opposing surface. The blue grasp requires a small (2.5 cm) grip aperture, and is thus optimal with respect to grasp aperture. The green grasp requires a large grip aperture (12.5 cm) and is thus sub-optimal. **(B)** For the two selected grasps in panel (A), we plot the optimality of the grasps (in normalized, arbitrary units) for each of the 4 optimality criteria, and the difference in optimality between grasps. **(C)** The difference in grasp optimality is shown for all pairs of grasps selected on all 16 objects, 4 per optimality criteria. Red indicates the selected near-optimal grasp is better than the selected sub-optimal grasp. Each column corresponds to one of the 16 objects employed in the study. The object and grasps in panel (A) correspond to the second column of the Aperture subplot in panel (C).

#### 2.3.2 Procedure

Experiment 1 consisted of a vision session followed by a grasping session. In each session, all objects were presented in random order. In a single trial of either session, participants were instructed to judge which of the two predefined grasps marked on the object was better. No specific definition of grasp quality was given to participants. In the vision session, no physical contact with the objects was allowed. Participants were instructed to imagine both grasp movements and verbally report which of the two grasps they thought was best. In the grasping session, participants executed both grasps and verbally reported which grasp was best. Participants were instructed to perform imagined and real grasps with a precision grip, i.e. using only thumb and index finger.

Prior to the experiment, participants were introduced to the objects. All stimuli were laid out on a table, the meaning of the stickers was explained, and participants were instructed to view (but not touch) the objects from all angles. Participants were familiarized with the weight of beech wood and brass by placing a wooden bar and a brass bar in sequence on the participants’ outstretched palm for a few seconds. Between trials of both sessions, and between grasps within one trial, we ensured that participants did not see the experimenter manipulating the objects by asking participants to keep their eyes closed until the objects were positioned.

In the vision session, once the stimulus was positioned at the starting location at its specific orientation, participants (with their head positioned on the chinrest) were instructed to open their eyes and visually explore the object. The experimenter then instructed the participant to imagine both blue and green grasps (in random order) and report which was best, with no time limit. During the vision session, participants were instructed to keep both hands on their thighs to prevent them from attempting pantomime grasps.

In the grasping session, on each trial participants positioned their head on the chinrest, and their thumb and index finger at the starting location. Once the stimulus was positioned, participants opened their eyes and the experimenter specified which grasp to attempt first (green or blue, in random order to minimize trial order effects; Maiello et al., 2018). Once the participant reported they were ready, an auditory cue specified the beginning of the grasping movement. Participants were required to reach, grasp, pick up and move the object onto the goal location, and return their hand to the starting position, all within three seconds. Prior to the second grasp, the experimenter positioned the current object back on its starting location while participants kept their eyes closed. Once the object was positioned, the procedure was repeated for the second grasp.

### 2.4 Experiments 2a and 2b

Experiment 2a was a replication of Experiment 1, except that we only employed a subset of the conditions and we recorded participants’ grasp movements during the grasping session using the ZED mini stereo camera. Compared to Experiment 2a, Experiment 2b contained an additional experimental session where participants evaluated grasp quality from the videos of participants from Experiment 2a.

#### 2.4.1 Stimuli

In Experiments 2a and 2b we employed only 6 objects out of the 16 employed in Experiment 1. This subset of conditions was selected so that participants would be at chance performance in the vision condition and significantly above chance in the grasping condition.

#### 2.4.2 Procedure

The procedure of Experiment 2a was identical to that of Experiment 1, except with fewer conditions.

In contrast to Experiment 1 and 2a, Experiment 2b consisted of three sessions: first a vision, then a video session, followed by a grasping session. The first (vision) and third (grasping) sessions were identical to the first and second sessions of Experiment 2a. In the video session of Experiment 2b, participants were shown videos of participants from Experiment 2a grasping the objects at the predefined grasp locations. Participants across Experiments 2a and 2b were yoked: each participant from Experiment 2b saw and evaluated the grasps from only one participant from Experiment 2a. The videos were taken from the left lens of the Zed mini stereo camera. Participants sat in front of a computer monitor.

On each trial, a dialogue box informed subjects which of the two grasps (green or blue) they would be viewing first. Participants started the video with a mouse click. Once the first grasp video was shown, a dialogue box informed participants they would be viewing the second grasp, and once again, participants started the video. Each video was shown only once. After participants had viewed both videos, they reported, via mouse click, which of the two grasps was better.

#### 2.5 Analyses

Data analysis was performed in Matlab version R2018a. Differences from chance performance and between group means were evaluated via unpaired and paired t-tests, as appropriate (p-values < .05 were considered statistically significant). We also report the 95% highest density interval (95% HDI) of the difference from chance or between group means, obtained via Bayesian estimation (Kruschke, 2013) using the Matlab Toolbox for Bayesian Estimation by Nils Winter. We compute effect size as 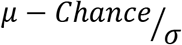 in case of differences from chance, and as 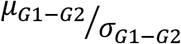 in case of differences between group means. As we are interested in fairly moderate effects (Cohen, 1988), we define a region of practical equivalence (ROPE) on effect size from −0.4 to 0.4. In cases where no statistically significant difference is observed using frequentist hypothesis testing, we use this ROPE to assess how credible the null hypothesis is that there exist no meaningful differences from chance or between group means (Kruschke, 2011). In such cases, we report the effect size and percentage of its posterior distribution that falls within the ROPE.

## 3 Results

### 3.1 Experiment 1: Participants can report whether grasps are optimal, and perform better when allowed to execute the grasps

In Experiment 1, we asked participants to perform imagined and real grasps on 16 objects and to report which of two predefined grasp locations was best. Figure 3A, shows that participants were significantly above chance at judging grasp optimality when using vision alone (t(20) = 6.63, p = 1.9*10^-06^; 95% HDI = [11, 22]) and also when physically executing the grasps (t(20) = 15.79, p = 9.3*10^-13^; 95% HDI = [25, 33]). Additionally, haptic and/or proprioceptive cues in the grasping session significantly improved participant’s judgements compared to the vision session (t(20) = 5.14, p = 5*10^-05^; 95% HDI = [8, 19]). Percent correct grasp optimality judgments for individual objects, grouped by optimality conditions, are shown in Supplementary Figures 1-4. Note that we do not compare performance across optimality conditions as we did not equate difficulty across conditions, and even within the same condition task difficulty and performance could vary markedly. Figure 3B further shows that participants who performed poorly in the vision session gained the most from physically executing the grasps: there was a strong, inverse relationship between grasping benefit^1^ and performance in the vision session (r = −0.73, p = 2*10^-4^).

**Figure 3.**
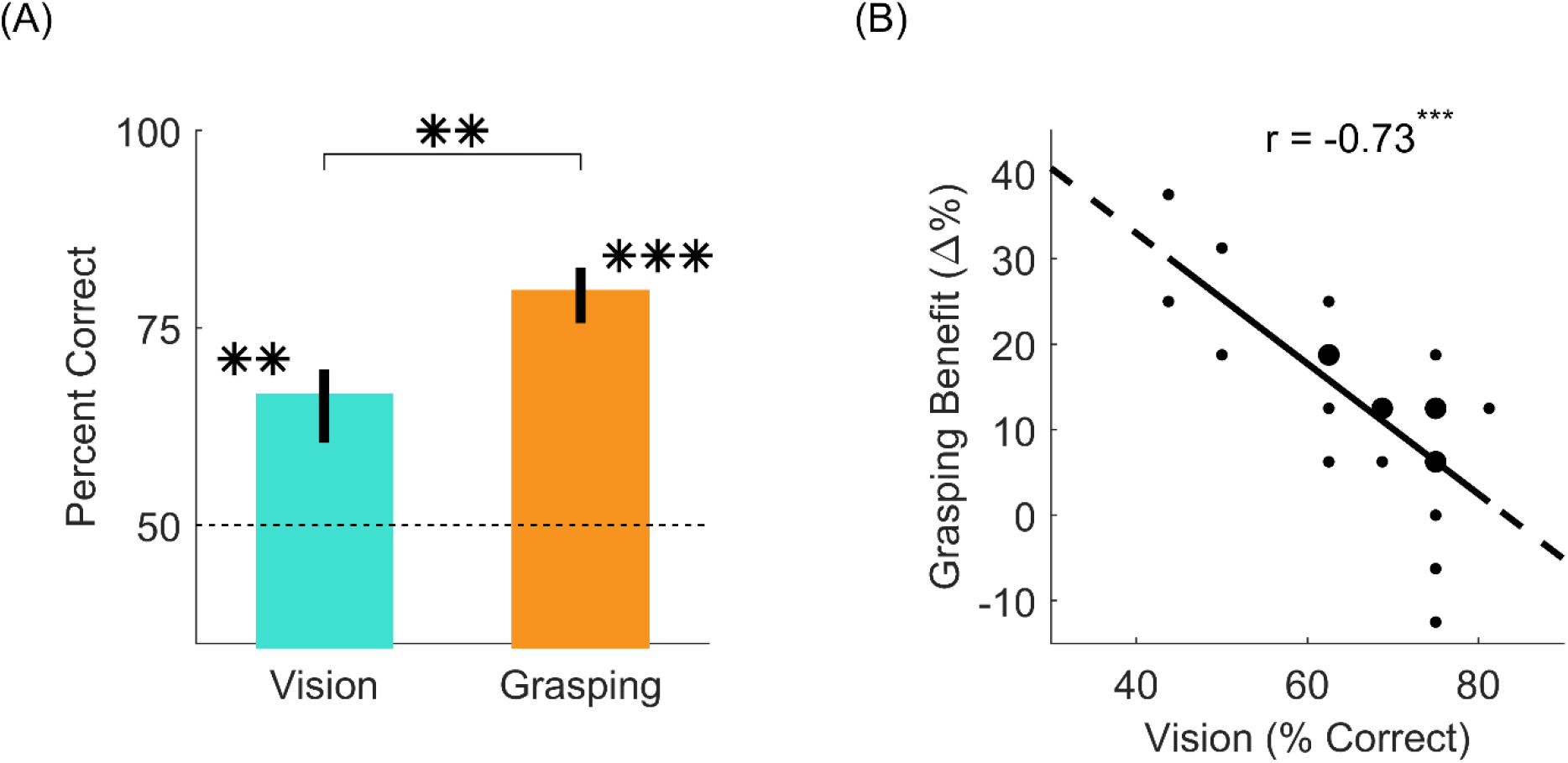
Judgments of grasp optimality using vision and grasping. **(A)** Percent correct grasp optimality judgments for the vision session (left), and the grasping session (right), averaged across objects and participants. Error bars indicate 95% bootstrapped confidence intervals of the mean. Chance performance is 50% correct (dotted line). **(B)** The grasping benefit (delta percent) as a function of the performance in the vision session, for each individual participant. The size of each dot represents the number of occurrences for each data point (one occurrence for small dots, two for large dots). Black line is best fitting linear regression line. **p<0.01; ***p<0.001.

### 3.2 Experiment 2: Visual and proprioceptive information *during grasping* are redundant for evaluating grasp optimality

The results from Experiment 1 suggest that participants are better at judging grasp quality when they perform the grasp. However, Experiment 1 leaves open whether the performance increase is due to the sensorimotor or visual feedback during grasp. In Experiment 2, we tested whether visual cues from real grasp movements were sufficient to improve grasp optimality judgements. In Experiment 1, performance varied across optimality criteria and individual objects.

Therefore, we selected the subset of conditions from Experiment 1 that showed the largest difference between the vision and grasping session. Figure 4A shows that for these conditions, participants were at chance in the vision session (t(20) = 0.5, p = 0.62; 95% HDI = [-8, 13], effect size = 0.11, 88% in ROPE), above chance when physically executing the grasps (t(20) = 10.25, p = 2.1*10^-09^; 95% HDI = [29, 40]), and performance in the grasping session was significantly improved compared to the vision session (t(20) = 4.81, p = 1.1*10^-4^; 95% HDI = [19, 46]).

**Figure 4.**
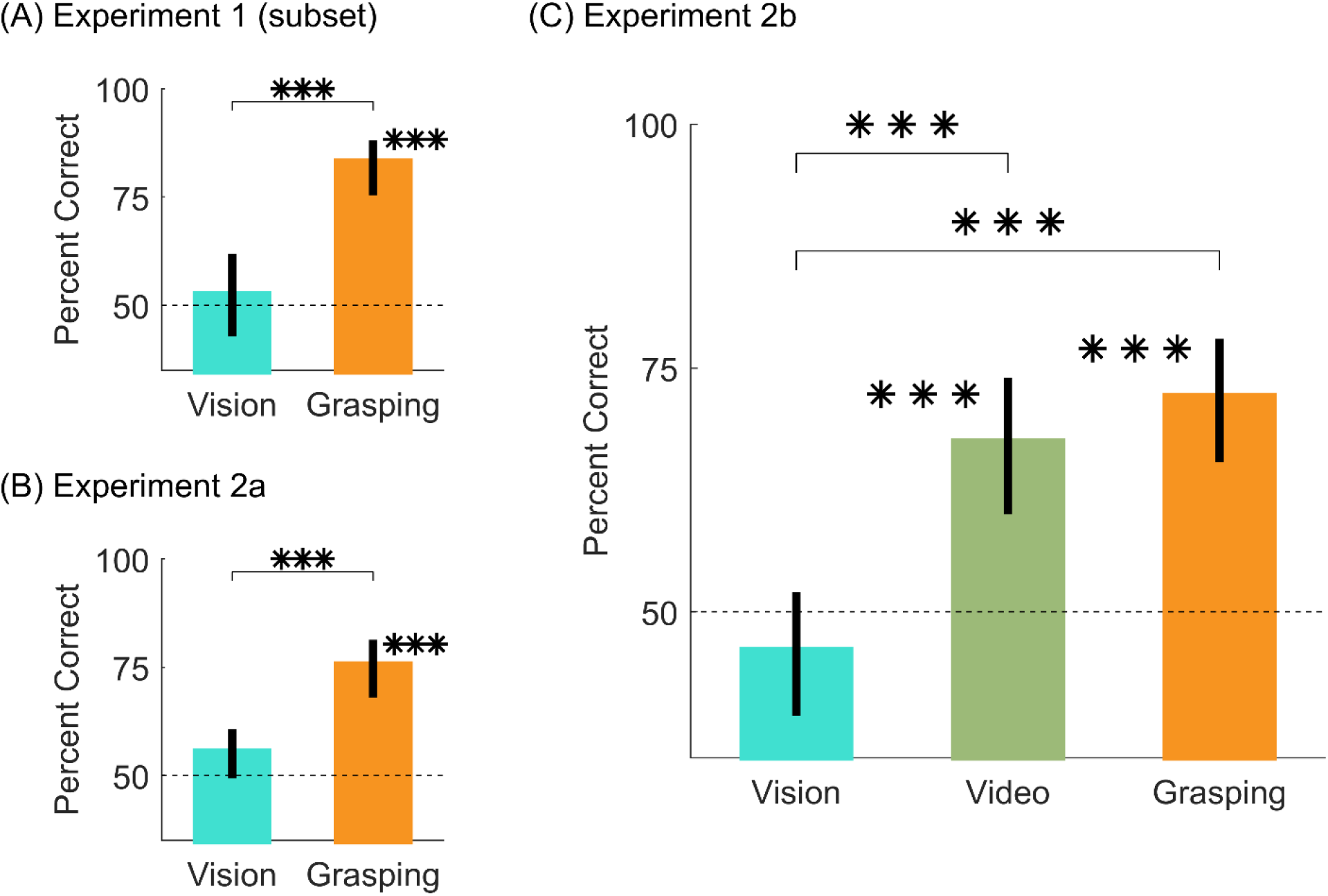
Results from Experiment 2. **(A,B)** Percent correct grasp optimality judgments for vision and grasping sessions, averaged across objects and participants, for **(A)** the subset of conditions from Experiment 1 that drives the difference between vision and grasping, and **(B)** the same subset of conditions replicated in Experiment 2a. **(C)** Percent correct grasp optimality judgments for vision, video, and grasping sessions, averaged across objects and participants, for Experiment 2b. In all panels, error bars are 95% bootstrapped confidence intervals of the mean and chance performance is 50% correct (grey dotted line). ***p<0.001.

In Experiment 2a we replicated the results from Experiment 1 on this subset of conditions (Figure 4B): participants were at chance in the vision session (t(24) = 1.88, p = 0.073; 95% HDI = [-1, 12], effect size = 0.38, 53% in ROPE), above chance when physically executing the grasps (t(24) = 7.27, p = 1.7*10^-07^; 95% HDI = [18, 33]), and performance in the grasping session was significantly improved compared to the vision session (t(24) = 3.51, p = 0.0018; 95% HDI = [8, 32]). During the grasping session of Experiment 2 we also recorded videos of the participants executing the grasps from approximately the participants’ viewpoint. Example videos are shown in the Supplementary Material.

In Experiment 2b, participants performed a vision, a video, and a grasping session on the same conditions employed in Experiment 2a. Critically, in the video condition participants judged grasp optimality on videos of participants from Experiment 2a grasping objects at optimal and sub-optimal locations.

Similarly to Experiment 2a, Figure 4C shows that in Experiment 2b, participants were at chance in the vision session (t(24) = −1.19, p = 0.25; 95% HDI = [−11, 4], effect size = −0.24, 81% in ROPE). Conversely, participants were significantly above chance in both the video (t(24) = 4.58, p = 1.2*10^-4^; 95% HDI = [10, 26]) and grasping sessions (t(24) = 6.41, p = 1.3*10^-06^; 95% HDI = [15, 29]). Compared to the vision session, performance was significantly improved in both the vision (t(24) = 4.23, p = 3*10^-04^, 95% HDI = [10, 32]) and grasping sessions (t(24) = 6.35, p = 1.4*10^-06^, 95% HDI = [17, 35]). Finally, performance in the video and grasping sessions was equivalent (t(24) = 0.92, p = 0.36; 95% HDI = [-6, 16], effect size = 0.18, 83% in ROPE). Percent correct grasp optimality judgments for individual objects and optimality conditions for both Experiments 2a and 2b are shown in Supplementary Figure S5.

## 4 Discussion

When grasping objects guided by vision, humans select finger contact points that are near-optimal according to several physics- and biomechanics-based constraints (Klein, Maiello et al., 2020; Kleinholdermann et al., 2013). Whether these constraints are explicitly computed in the brain is unknown. Here, we demonstrate that humans can explicitly judge which of two potential grasps on an object is best, based on each of these constraints.

In our study, participants could distinguish near-optimal from sub-optimal grasp locations using vision alone, i.e. without physically executing grasps, presumably using motion imagery. This well aligns with the notion that motor imagery, the mental simulation of a motor task, relies on similar neural substrates as action planning and execution. For example, it is well established that simulated actions take the same time as executed ones (Decety et al., 1989; Jeannerod, 1995). This temporal similarity has also been shown in a task akin to the current study. Frak et al. (2001) asked participants to determine whether contact points marked on a cylindrical object placed at different orientations would lead to easy, difficult, or impossible grasps, without grasping the object. The time to make these estimates varied with object orientation and task difficulty, and closely matched the time taken to perform the grasps. These temporal matches hint that imagined and real actions might rely on similar neural computations. Indeed it has been shown that motor imagery recruits many of the same visuomotor areas of the brain, from early visual cortex (Monaco et al., 2020; Pilgramm et al., 2016; Zabicki et al., 2016), throughout the dorsal stream and the parietal lobe leading to primary motor cortex M1 (Hétu et al., 2013), that are directly involved in action planning and execution (Hardwick et al., 2018).

In Experiment 1 of our study, judgements of grasp optimality improved when participants were required to execute the grasps, and this improvement was strongest in participants who performed poorly using vision alone. What drove this improvement? In the grasping sessions, participants were asked to grasp, lift and place the object at a goal location within three seconds. However, they had unlimited time to plan the grasps prior to each trial. The planning stage in the grasping sessions was thus similar to the vision sessions. Therefore, in both sessions participants could build hypotheses about which grasp should be easier to execute, but only in the grasping sessions could they test these hypotheses against their own sensorimotor feedback. Specifically, if participants needed to make corrective changes once a movement had been initiated, it is possible that the difference between this event and the original motor intention could have reached consciousness and improved their judgements. However, previous research has shown that the recalibration of reach-to-grasp movements through haptic feedback occurs outside of perceptual awareness (Mon-Williams & Bingham, 2007). If participants could not consciously access the corrections to their original motor plans, crucial clues to indicate that a grasp was sub-optimal could be provided by tactile feedback from object slippage (Johansson & Westling, 1984), the need to apply greater grip forces than anticipated (Lukos et al., 2013), or proprioceptive feedback indicating awkward joint configurations (Rosenbaum et al., 2001).

Tactile and proprioceptive feedback were not the only sources of information that could have aided judgements in the grasping session. Participants could also visually assess the characteristics of the own movements, such as the speed and trajectory of the limb. These sources of visual information are known to play a strong role in grasp execution, as removing them changes the kinematics of grasping movements (Connolly & Goodale, 1999). Additionally, even if visual information from object roll during grasps does not influence the calibration of digit placement and force control (Lukos et al., 2013), lifting without visual feedback does impair fingertip force adaptation (Buckingham et al., 2011; Buckingham & Goodale, 2010). We therefore wondered whether these sources of visual information alone could aid judgements of grasp optimality.

In Experiment 2, we indeed found that viewing videos of other participants grasping near-optimal and sub-optimal grasps was sufficient for observers to reach the same level of performance at reporting which grasp was best as when actually executing grasps. This does not mean that in the grasping sessions participants did not rely on tactile and proprioceptive feedback. It suggests instead, that visual and tactile/proprioceptive feedback may be redundant with visual information in evaluating grasp quality. This could help explain how humans are able to exploit action observation more generally. For example, humans are able to acquire useful information, such as object weight, by simply observing the movement kinematics of others (Bingham, 1987; Hamilton et al., 2007). Additionally, observing others execute grasping tasks, particularly when they make errors, can improve one’s own grasping performance (Buckingham et al., 2014). Observing one’s own grasps, particularly when making errors, could thus link visual and tactile/proprioceptive information about grasp quality. This in turn would allow us to learn how best to grasp a novel object by simply looking at someone else grasping it.

### 4.1 Limitations and Future directions

Our findings reinforce the notion that motor imagery and action observation play an important role in learning complex motor tasks (Gatti et al., 2013). For this reason, motor imagery and action observation have also shown promise in aiding and strengthening motor rehabilitation techniques in a variety of neurological conditions (de Lange et al., 2008; Malouin et al., 2013; Mateo et al., 2015; Mulder, 2007; Sharma et al., 2006; Zimmermann-Schlatter et al., 2008). Within this context, our model-driven method of selecting optimal—and particularly sub-optimal—grasps could be used to guide and strengthen mental imagery and action observation techniques for motor rehabilitation. For example, patients could be made to imagine, observe, and execute grasps to object locations, selected through our modeling approach, which contain the most useful information for re-learning grasping movements.

Even in the grasping sessions however, in about 20% of trials participants did not agree with the model predictions. Does this mean participants could not access the information about grasp quality? We believe it is more likely that the model predictions are incomplete. For example, the model does not take into account that for some grasps with high torques, the objects might rotate and come to rest against a participants’ palm, stabilizing an otherwise potentially unstable grasp. Additionally, in the current work we did not account for the different importance given by individual participants to the different constraints (Klein, Maiello et al., 2020). Inspect for example the data from the last panel of Supplementary Figure 2. Even though the selected sub-optimal grasp has much larger grasp aperture than the selected near-optimal grasp, the sub-optimal grasp has marginally less torque. Thus, if some participants gave much greater importance to the torque constraint, this might explain why their responses disagreed with model predictions.

The videos from Experiment 2 could provide some further insight into which visual cues participants were exploiting to determine grasp optimality during action observation. For example, in Supplementary Video 1 an observer might notice the different time it takes the participant to lift the same object with two different grasps, or the slight wobbling of the object when grasped in the uncomfortable hand orientation. In Supplementary Video 2, a prominent visual cue comes from the initial failure in computing a successful trajectory to the sub-optimal grasp. A quantitative analysis of the grasping kinematics contained in these videos, using for example novel image based tracking algorithms (Mathis et al., 2018), may reveal the exact nature of the visual information human participants exploit during action observation. The full video dataset from Experiment 2, as well as all other data from the study, are made freely available through the Zenodo repository (doi: xx.xxxx/zenodo.xxxxxxx upon publication).

Finally, our approach could be further developed to investigate the neural underpinning of visual grasp selection. The current study demonstrates how, through the computational framework described in (Klein, Maiello et al., 2020), we can identify grasps on arbitrary objects that isolate the individual components of grasp selection. In future studies, these unique grasp configurations could be employed as stimuli for targeted investigations of brain activity, making it possible to pinpoint the neural loci of each of the visuomotor computations underlying grasp planning and execution.

### 4.2 Conclusion

We show that humans are capable of judging the relative optimality between different possible grasps on an object. Humans can perform these judgments using vision alone, and can refine their estimates of grasp quality using visual and proprioceptive feedback during grasp execution. These abilities are likely a key component of how humans visually select grasps on objects. Remaining challenges will be to identify where and how grasp optimality is learned and computed in the brain in order to guide grasp planning and execution.

## 6 Author Contributions

GM, MS, LKK, VCP and RWF conceived and designed the study. GM and MS collected the data. GM analyzed the data. All authors wrote the manuscript.

## 8 Acknowledgments

This article is based on Marcel Schepko’s master’s thesis.

## 10 Supplementary Material

**Supplementary Figure 1.** Percent correct grasp optimality judgments, computed across participants from Experiment 1, for the 4 individual objects in the natural grasp axis conditions. In each panel, the top object demonstrates the approximate viewpoint of a participant. Thumb locations for selected grasps were marked on the objects in green or blue. The position of the opposing index finger was marked in yellow. The color code only served to mark and identify the grasps for participants, and was purposely unrelated to the grasp optimality. The middle and bottom object show the near-optimal and sub-optimal grasps respectively, with the objects rotated solely for illustrative purposes, to better show the selected grasp locations.

**Supplementary Figure 2.** As Supplementary Figure 1, except for the 4 individual objects in the grasp aperture conditions.

**Supplementary Figure 3.** As Supplementary Figure 1, except for the 4 individual objects in the minimum torque conditions.

**Supplementary Figure 4.** As Supplementary Figure 1, except for the 4 individual objects in the object visibility conditions.

**Supplementary Figure 5.** As Supplementary Figures 1-4, except for the 6 individual objects employed in Experiments 2a and 2b.

**Supplementary Video 1.** Representative participant from Experiment 2a executing near-optimal (left) and sub-optimal (right) grasps for one object belonging to the natural grasp axis conditions.

**Supplementary Video 2.** Representative participant from Experiment 2a executing near-optimal (left) and sub-optimal (right) grasps for one object belonging to the object visibility conditions.

## 11 Data Availability Statement

The datasets generated and analyzed for this study will be made available from the Zenodo repository upon publication (doi: xx.xxxx/zenodo.xxxxxxx).

## Footnotes

1 Grasping benefit was defined as: %*Correct_Grasping_* – %*Correct_Vision_*

