## Supplementary Figures for "Humans can visually judge grasp quality and refine their judgments through visual and haptic feedback"

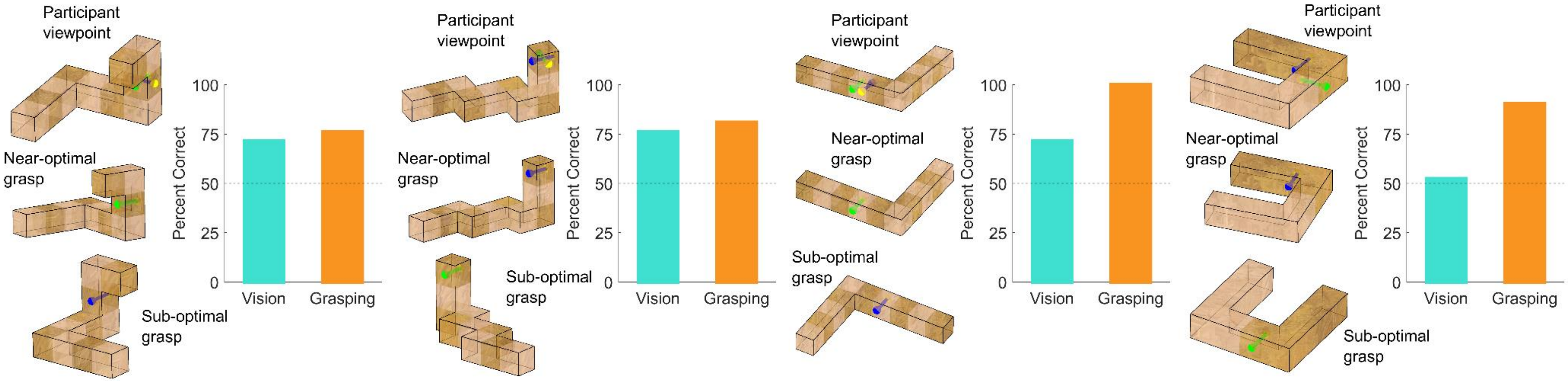


**Supplementary Figure 1.** Percent correct grasp optimality judgments, computed across participants from Experiment 1, for the 4 individual objects in the natural grasp axis conditions. In each panel, the top object demonstrates the approximate viewpoint of a participant. Thumb locations for selected grasps were marked on the objects in green or blue. The position of the opposing index finger was marked in yellow. The color code only served to mark and identify the grasps for participants, and was purposely unrelated to the grasp optimality. The middle and bottom object show the near-optimal and sub-optimal grasps respectively, with the objects rotated solely for illustrative purposes, to better show the selected grasp locations.

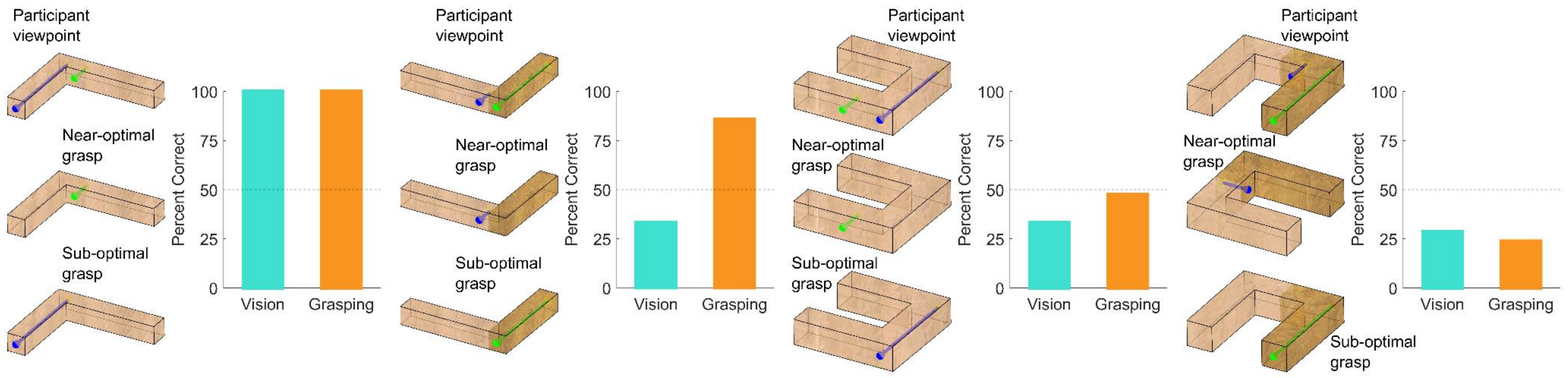


**Supplementary Figure 2.** As Supplementary Figure 1, except for the 4 individual objects in the grasp aperture conditions.

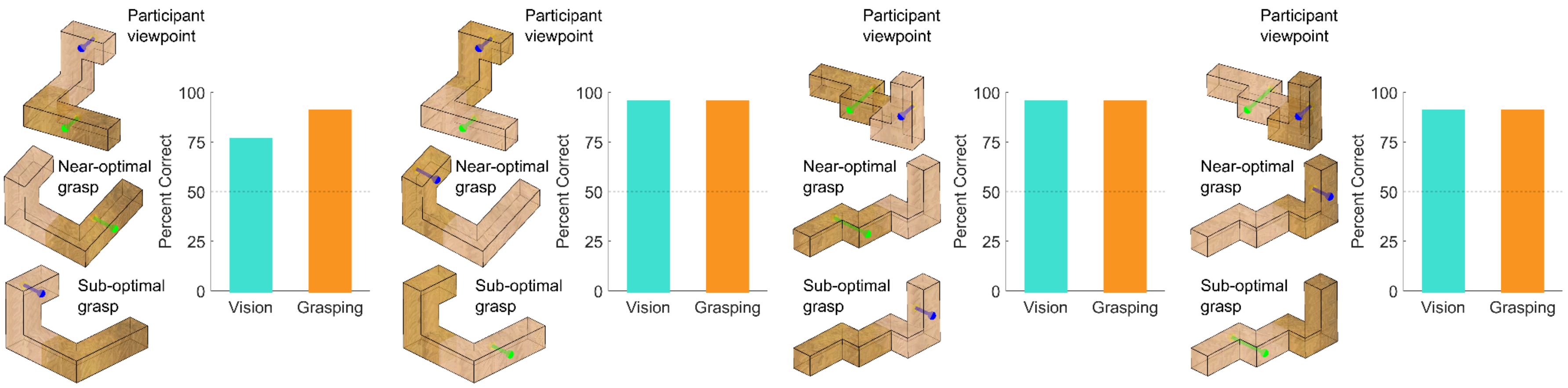


**Supplementary Figure 3.** As Supplementary Figure 1, except for the 4 individual objects in the minimum torque conditions.

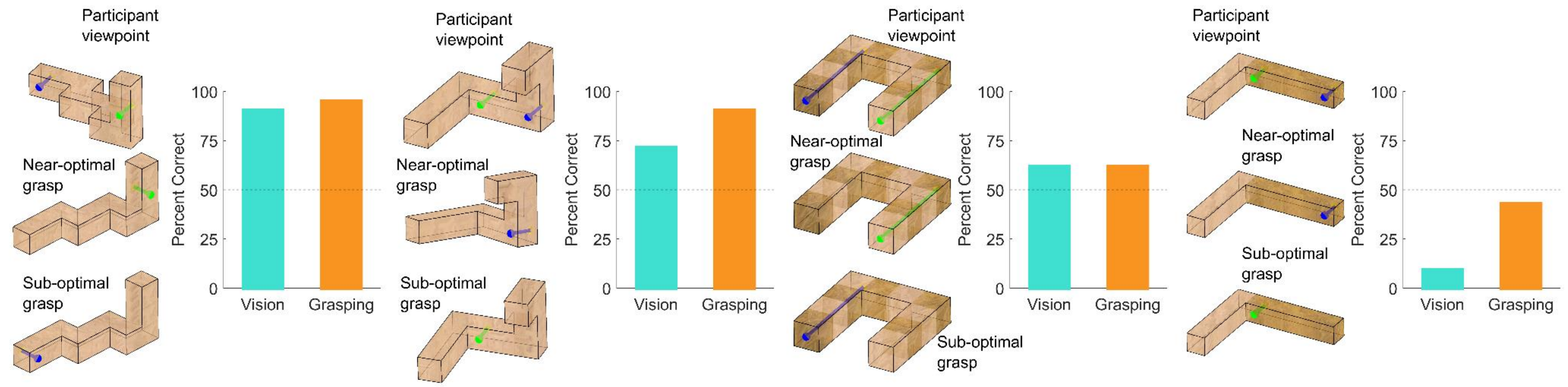


**Supplementary Figure 4.** As Supplementary Figure 1, except for the 4 individual objects in the object visibility conditions.

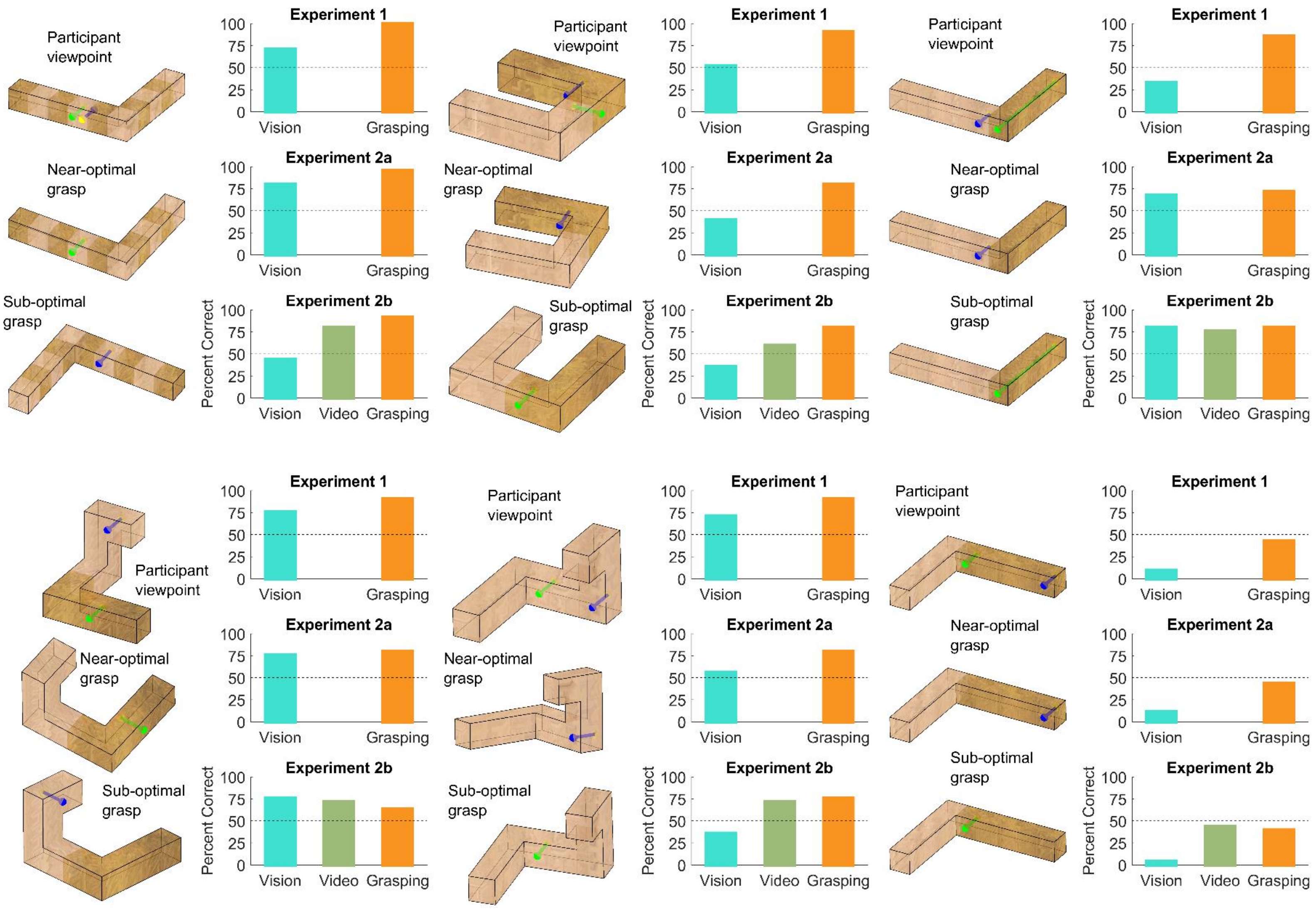


**Supplementary Figure 5.** As Supplementary Figures 1-4, except for the 6 individual objects employed in Experiments 2a and 2b.
